# Snapshot of Defense Systems in Multidrug Resistant *Klebsiella pneumoniae*

**DOI:** 10.1101/2025.04.16.649094

**Authors:** Tosin Y Senbadejo, Samuel O. Ntiamoah, Abiola Isawumi

## Abstract

**Objectives:** The defense mechanisms in bacterial pathogens protect them from host immune systems, bacteriophage infections and stringent environmental conditions. This study explores the defense-systems in multidrug resistant *Klebsiella pneumoniae* isolated from Ghanaian hospital ICUs focusing on CRISPR-cas, restriction-modification and toxin-antitoxin systems (TAs).

**Method:** Genomic DNA of *K. pneumoniae* environmental (NS2) and clinical (PS4) strains were subjected to whole genome sequencing using Illumina and assembled with SPAdes (v3.13.1). CRISPR-cas, restriction-modification and TAs were identified using PADLOC, defense finder and TADB3.0 respectively.

**Results:** The strains harbor diverse defense systems. Relative to reference *K. pneumoniae* with 10 defense systems, NS2 has twelve and PS4, five. CRISPR-Cas systems were found only in NS2, while both strains have type I, II and IV restriction and modification systems. The strains have > 30 characterized and novel TAs (type I, II, IV, VIII) similar to reference *K. pneumoniae*. NS2 harbors more TAs than PS4 both on chromosomes and plasmids. The strains have comparable resistance determinants to more than six classes of antibiotics.

**Conclusion:** The genome of strains encodes similar clinically relevant defense-systems indicating possibility of microbial exchange from fomites and humans. They could leverage the defense-systems to propagate resistance in high-risk environments such as the hospital. Fomite-resident strain with high levels of resistance could increase infection risk in the ICU; hence, should also be prioritized.

## Introduction

*Klebsiella pneumoniae* is an opportunistic pathogen implicated in hospital-acquired and community associated infections (Liang et al., 2022). As multidrug resistant pathogen, it contributes to public health challenges with consequent 22% to 72% annual mortality and morbidity rate (Poerio et al., 2022). Clinical and environmental *K. pneumoniae* shares antibiotic resistance (AMR) phenotypes, virulence and resistance genes (Rocha et al., 2022). Its severity and invasiveness is attributed to genome complexity and acquisition of factors facilitating persistence and evasion of host immune systems (de Sales et al., 2022). Some of these genomic factors are defense systems that play relevant roles in virulence and spread of multidrug resistance.

CRISPR-Cas, TAs, and restriction-modification (R-M) are defense systems involve in bacterial colonization, virulence regulation, biofilm formation, persistence, spread of genetic elements, and immunity against phages (Horesh et al., 2020). CRISPR-Cas systems in *K. pneumoniae* interfere with acquisition of phages and genetic elements that harbor AMR genes; therefore reduce AMR levels (Li et al., 2018). It has also been indicated that they can be explored for development of novel antimicrobial agents in the treatment of MDR *K. pneumoniae* infections (Xu et al., 2019).

Restriction and modification systems protect host bacterial cells from viruses and other bacteria (Bogdanova et al., 2008). R-M system is the most abundant and studied defense system in prokaryotes (Tesson et al., 2022). Type II R-M is the most abundant in bacterial genomes and in *K. pneumoniae*, KpnAI, KpnBI, Kpn2I are the most common (Chin, 2000). They provide the first line of immunity with restriction endonuclease (R) and methyltransferase (M). The R recognizes foreign DNA sequence and introduces double-stranded breaks around the recognition site while the M recognizes host DNA sequence and methylates to prevent cleavage and site recognition by R (Bogdanova et al., 2008). The type II systems are mostly plasmid-encoded for horizontal gene transfer (Rodic et al., 2017) and ensure bacterial persistence.

Toxin-antitoxin systems contribute to bacterial cell growth arrest and survival (Li et al., 2023). Also, it helps in genetic material maintenance, biofilm formation, resistance to stresses, virulence and pathogenesis (Kamruzzaman et al., 2021). TAs comprises a stable toxin and a labile antitoxin that can either be protein or RNA depending on the type. There are eight chromosomal or plasmid-mediated TAs (Type I -VIII) within bacterial genome (Jurėnas et al., 2022). In Ghana, there is paucity of data on defense systems in MDR *K. pneumoniae*, especially those isolated from Intensive Care Unit (ICU). It is therefore important to profile defense systems in the newly emerging MDR clinical and environmental *K. pneumoniae*, as AMR is increasingly becoming a burden in Ghana.

## Materials and Methods

### Isolation, characterization and annotation of the bacterial strains

*Klebsiella pneumoniae* (NS2 and PS4) strains were obtained from ABISA Bacterial Culture Library. They were isolated from septicemia patients (PS4) and hospital ICU fomites (NS2). The gDNA was extracted using QIAGEN bacterial-DNA kit, sequencing via Illumina platform, and QC of raw reads using FastQC. Prokka (v1.14.6) and RAST were used for gene prediction and annotation of resulting contigs from spade assembler. Average Nucleotide Identity (ANI) was determined with JSpecies-WS. The nucleotide and protein sequences of the coding genes were used as data files for profiling the defense systems. The reference *K. pneumoniae* (HS11286) was downloaded from the NCBI database.

### The defense systems, plasmids and resistance genes

Prokaryotic Antiviral Defense Locator (PADLOC) was used for profiling of defense systems. The genes encoding the defense systems were identified by matching protein sequences with curated database (Hidden Markov Models-HMM) with >700 families of defense systems related proteins. Defensefinder and KEGG were used for confirmation. The PADLOC output was loaded into Geneious prime for visualization of CRISPR-Cas and R-M systems. TADB3.0 was used to predict different types of TA pairs. Existing sequences of experimentally validated TA-proteins were also inputted into TBLASTN and PSI-BLAST programs. Searches were performed against the annotated MDR *K. pneumoniae* genomes and plasmids with E-value >0.05. Mobile genetic element finder (MGE-Finder v1.0.3) was employed for identifying plasmids (and other MGE). Comprehensive Antibiotic Resistance Database (CARD) and Resistance finder (ResFinder) for resistance genes.

## Results and Discussion

### General Features of multidrug resistant *Klebsiella pneumoniae*

Based on the genome sequencing, NS2 has 5,713,857 read sequences in 253 contigs with a GC content (57.1%), 5751 coding sequences and 94 RNAs while PS4 has 5,522,745 sequences in 126 contigs with a GC content (57.3), 5413 coding sequences and 93 RNAs (**Table 1**). The strains have similar features with annotated reference *K. pneumoniae*. They have ANI values of 98 – 99% with those from database based on BLAST and MUMmer results (Supplementary). NS2 as an environmental strain has a larger genome, more plasmids and hypothetical proteins than PS4. Plasmids harbor genes that encode resistance and virulence (supplementary). It has been indicated that *K. pneumoniae* resident on fomites (medical devices) harbor resistance markers, plasmids and stress-related genes (Rocha et al., 2022). The genome data has been submitted to NCBI genome database with **SUB14552324** designation and will be made available on request.

**Table 1:** Comparative genome features of *Klebsiella pneumoniae* strains.

| Parameters | NS2 (Environmental strain) | PS4 (clinical strain) |
| --- | --- | --- |
| Sequence size | 5713857 | 5522745 |
| Number of Contigs | 253 | 126 |
| Longest contig size | 428242 | 992492 |
| GC content (%) | 57.1 | 57.3 |
| Genes | 5343 | 5506 |
| Protein-coding genes | 5249 | 5079 |
| Number of Plasmids | 4 | 1 |

### Defense systems identification

The genomes of the strains contain chromosome and plasmid-borne Type I, II and IV R-M systems. They are also abundant in the ‘pathogenicity islands’ indicating possible roles in pathogenicity and virulence. CRISPR-Cas systems including CRISPR-CAs Type I and subtype I-E were found in NS2 environmental isolate (**Figure 1**). In similar study, Type I-E and subtype I-E were dominant in clinical isolates in association with higher virulence, acquisition of plasmids, other MGEs and MDR genes in *K. pneumoniae* (Alkompoz et al., 2023). The clinical isolate (PS4) carried MGEs encoding genes that could confer AMR, heavy metals tolerance and stress adaptation. In comparison with reference strain, NS2 and PS4 have 36 genes (14 systems) and 17 genes (6 systems) respectively. This might indicate acquisition of extra defense systems from the environment or through selection pressure, especially at the ICU with significant reliance on antibiotics. CRISPR-Cas and R-M systems contribute to evolution and interactions between MGEs and bacterial hosts (Koonin & Makarova, 2017). There are more than thirty characterized and novel TAs. Three of the four NS2 plasmids have TAs (Supplementary). The frequency of Type II TAs are higher (**Table 2**), similar to previous study (Sberro et al., 2013). Majority of the TAs were localized in the chromosomes, contrary to what has been reported (Leplae et al., 2011). Also, some of the TAs were on the genomic island alongside R-M systems and other virulence genes.

**Table 2:**
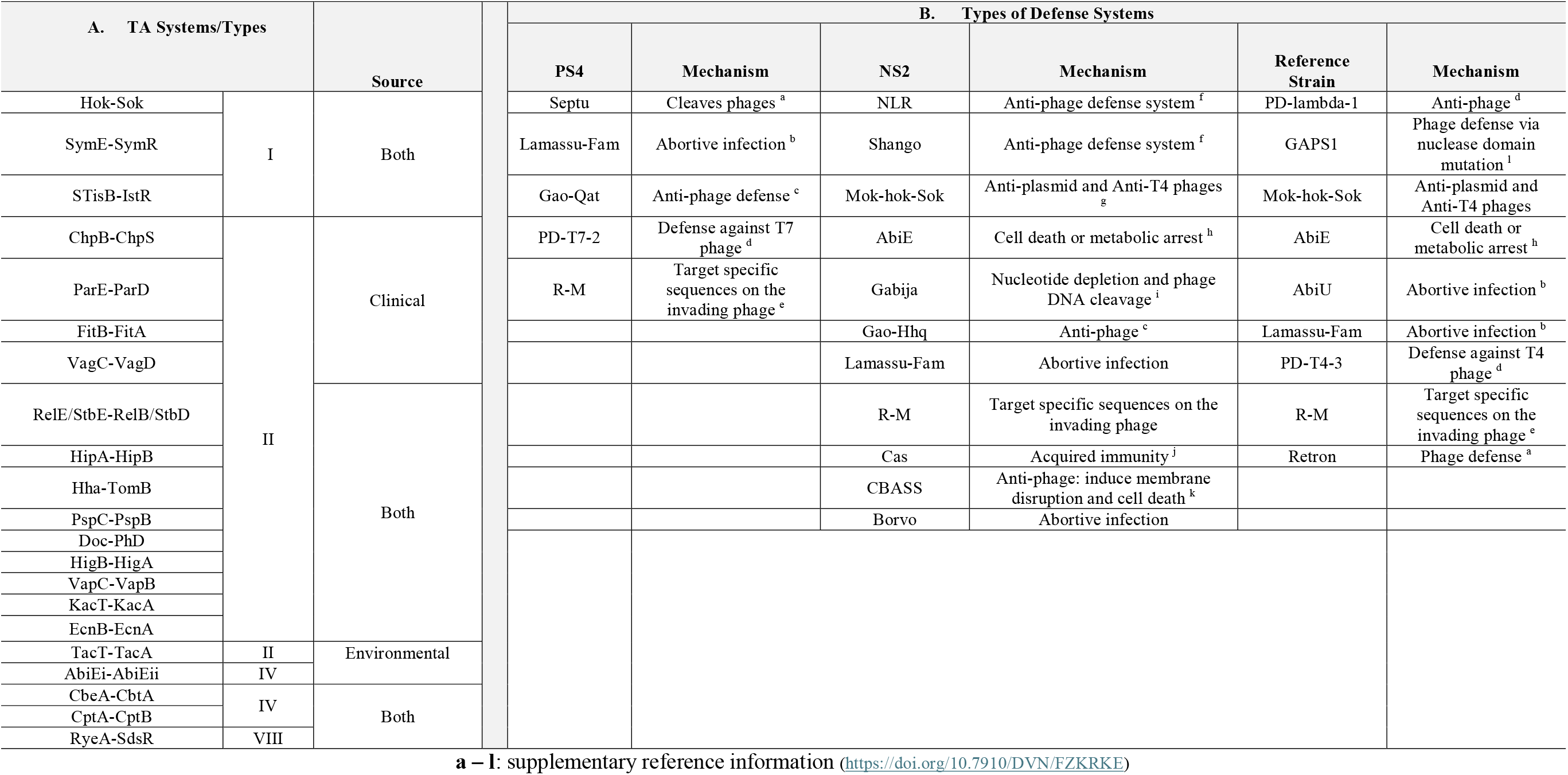
Defense Systems in Clinical and Environmental *Klebsiella pneumoniae* strains.

**Figure 1:**
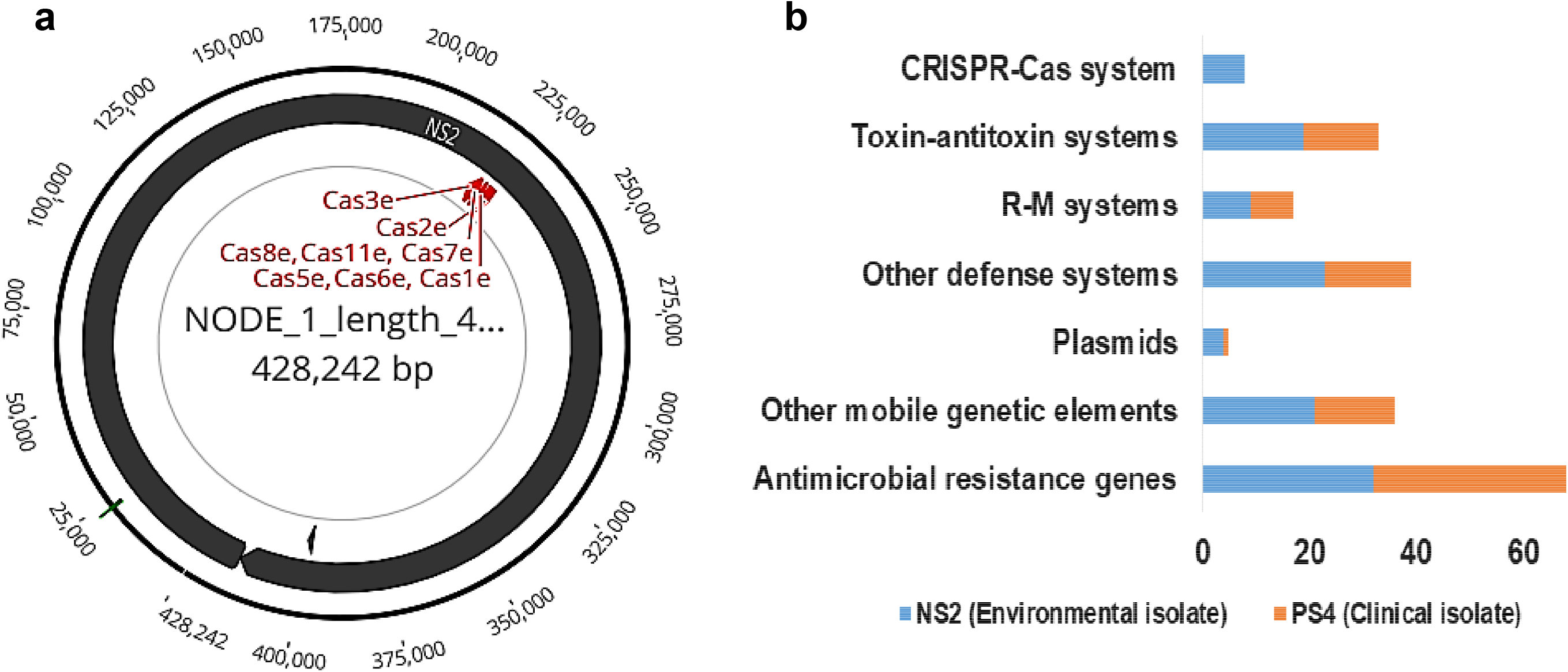
Genome profiles of Klebsiella pneumoniae (a) CRISPR-Cas identified in clinical isolate (NS2); (b) Defense systems and other genome features.

## Conclusion

This is the first study to characterize CRISPR-cas, R-M and TAs in environmental and clinical *K. pneumoniae* strains from Ghanaian hospital ICUs. The strains have clinically relevant and comparable defense systems, thus indicating possibility of fomites and humans’ microbial exchange. This can also promote the spread of AMR, virulence genes and increase the risk of hospital acquired infections. The high levels of TAs and R-M in the genome of the strains might increase the expression of MGEs, AMR virulence genes and facilitate bacterial persistence under stress. These defense systems could serve as tools for prevention and control of MDR *K. pneumoniae* considering their abundance and contribution to virulence and pathogenicity.

## Acknowledgements

The authors would like to acknowledge members and staff of AMR Research Group (led by Dr Abiola Isawumi) at West African Centre for Cell Biology of Infectious Pathogens (WACCBIP), Department of Biochemistry, Cell and Molecular Biology, University of Ghana for technical support.

## Data Availability Statement

All data generated and analysed during this study are included in this published article (and its supplementary information files – Isawumi, Abiola, 2024, “Defense Systems in *Klebsiella pneumoniae*”, https://doi.org/10.7910/DVN/FZKRKE, Harvard Dataverse, V2)

## LIST OF ABBREVIATIONS

NS2: Environmental *Klebsiella pneumoniae* strain
PS4: Clinical *Klebsiella pneumoniae* strain
gDNA: Genomic DNA
TA: Toxin-antitoxin systems
PADLOC: Prokaryotic Antiviral Defense Locator
CRISPR: Clustered Regularly Interspaced Short Palindromic Repeats
R-M: Restriction modification systems
R: Restriction endonuclease M-Methyltransferase
ICU: Intensive care unit
AMR: Antimicrobial resistance
RAST: Rapid Annotation Subsystem Technology
ANI: Average Nucleotide Identity
MGE: Mobile Genetic Element
CARD: Comprehensive Antibiotic Resistance Database
ResFinder: Resistance Finder
MDR: Multidrug resistant
BLAST: Basic Local Alignment Sequencing Tool
PSI-BLAST: Position-Specific-Iterated BLAST

