## Supplementary for "Snapshot of Defense Systems in Multidrug Resistant *Klebsiella pneumoniae*"

**NS2 - PLASMIDFINDER**

| **Plasmid** | **Identity** | **Query / Template length** | | **Contig** | **Position in contig** | **Accession number** |
| --- | --- | --- | --- | --- | --- | --- |
| IncFIA(HI1) | 98.45 | | 387 / 388 | NODE_34_length_23730_cov_52.7371 | 2495..2881 | [AF250878](http://www.ncbi.nlm.nih.gov/nuccore/AF250878) |
| IncFIB(K) | 100 | | 560 / 560 | NODE_40_length_12559_cov_84.0491 | 4153..4712 | [JN233704](http://www.ncbi.nlm.nih.gov/nuccore/JN233704) |
| IncFII(K) | 97.3 | | 148 / 148 | NODE_22_length_75109_cov_91.7874 | 9121..9268 | [CP000648](http://www.ncbi.nlm.nih.gov/nuccore/CP000648) |
| IncR | 100 | | 251 / 251 | NODE_37_length_15203_cov_56.898 | 9403..9653 | [DQ44957](http://www.ncbi.nlm.nih.gov/nuccore/DQ449578) |

**PLASMIDS HARBOUR RESISTANCE AND VIRULENCE GENES**

| NS2 | Resistance genes (RESFINDER) | Virulence genes (VFDB) | Toxin-Antitoxin systems (TADB2.0) |
| --- | --- | --- | --- |
| Plasmids | *blaTEM-1B, qacE, OqxB, OqxA, aph(3’)-lb, aph(6)-ld, fosA, blaSHV-65, catA1* | *fimH, mrkA, fyuA, irp2, iutA, clpK1* | *TacT/TacA, HigB/HigA,Hok/Sok, Hha, hipB, relB,* |

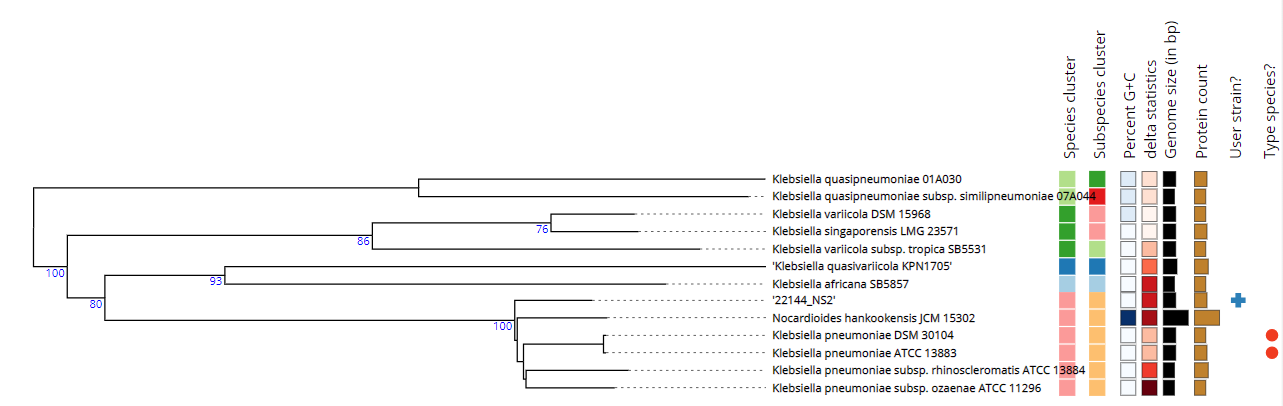

**NS2 - TYPE STRAIN GENOME SERVER (TYGS) PHYLOGENY BASED ON WHOLE GENOME SEQUENCE**

**PS4 - PLASMIDFINDER**

| **Plasmid** | **Identity** | **Query / Template length** | **Contig** | **Position in contig** | **Accession number** |
| --- | --- | --- | --- | --- | --- |
| IncFIB(K)(pCAV1099-114) | 99.64 | 560 / 560 | NODE_17_length_83010_cov_58.2909 | 73877..74436 | [CP011596](http://www.ncbi.nlm.nih.gov/nuccore/CP011596) |

PLASMID HARBOUR RESISTANCE AND VIRULENCE GENES

| PS4 | Resistance genes (RESFINDER) | Virulence genes (VFDB) | Toxin-Antitoxin systems (TADB2.0) |
| --- | --- | --- | --- |
| Plasmid | *blaSHV-85, blaSHV-40, blaTEM-IB, Sul2, OqxB, OqxA, aph(6)-ld, fosA, blaSCO-1, aac(3)-lla, dfrA14, blaCTX-M-15.* | *fimH, mrkA, fyuA, irp2, iutA, clpK, traT, nlpI* | *hepT/mntA, tacA/tact, vapC, relB, Hha* |

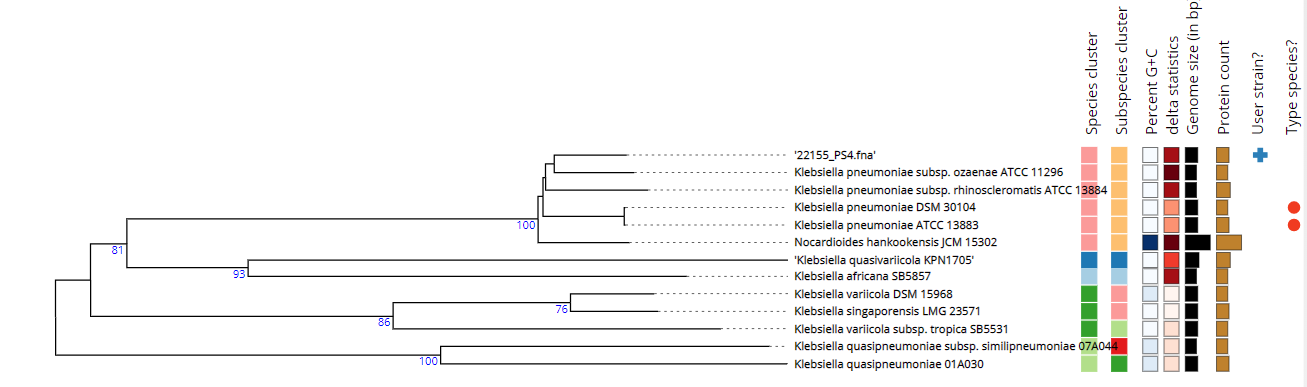

**PS4 - TYPE STRAIN GENOME SERVER (TYGS) PHYLOGENY BASED ON WHOLE GENOME SEQUENCE**

|  | **22144 NS2.fasta** | **22155 PS4.fasta** | **Klebsiella pneumoniae 140 1040** | **Klebsiella pneumoniae 160 1080** | **Klebsiella pneumoniae KP ST11 OXA48** | **Klebsiella pneumoniae 440 1540** | **Klebsiella pneumoniae 646 1568** |
| --- | --- | --- | --- | --- | --- | --- | --- |
| **22144 NS2.fasta** | * | **99.22** | **99.25** | **99.25** | **99.27** | **99.25** | **99.26** |
| **22155 PS4.fasta** | **99.22** | * | **99.21** | **99.23** | **99.22** | **99.21** | **99.23** |
| **Klebsiella pneumoniae 140 1040** | **99.25** | **99.21** | * | **99.98** | **99.74** | **99.99** | **99.99** |
| **Klebsiella pneumoniae 160 1080** | **99.26** | **99.23** | **99.98** | * | **99.75** | **99.98** | **100** |
| **Klebsiella pneumoniae KP ST11 OXA48** | **99.27** | **99.22** | **99.74** | **99.75** | * | **99.74** | **99.75** |
| **Klebsiella pneumoniae 440 1540** | **99.25** | **99.21** | **99.99** | **99.98** | **99.74** | * | **99.98** |
| **Klebsiella pneumoniae 646 1568** | **99.26** | **99.24** | **99.99** | **100** | **99.75** | **99.98** | * |

**Average Nucleotide Identity (ANI) determined using JSpecies-WS**

|  | **22144 NS2.fasta** | **22155 PS4.fasta** | **Klebsiella pneumoniae 140 1040** | **Klebsiella pneumoniae 160 1080** | **Klebsiella pneumoniae KP ST11 OXA48** | **Klebsiella pneumoniae 440 1540** | **Klebsiella pneumoniae 646 1568** |
| --- | --- | --- | --- | --- | --- | --- | --- |
| **22144 NS2.fasta** | * | **99.22** | **99.25** | **99.25** | **99.27** | **99.25** | **99.26** |
|  |  | *-87.38* | *-87.3* | *-86.34* | *-89.93* | *-87.17* | *-86.39* |
| **22155 PS4.fasta** | **99.22** | * | **99.21** | **99.23** | **99.22** | **99.21** | **99.23** |
|  | *-90.57* |  | *-88.1* | *-87.55* | *-90.29* | *-87.91* | *-87.59* |
| **Klebsiella pneumoniae 140 1040** | **99.25** | **99.21** | * | **99.98** | **99.74** | **99.99** | **99.99** |
|  | *-90.37* | *-87.96* |  | *-97.89* | *-95.63* | *-99.68* | *-98.11* |
| **Klebsiella pneumoniae 160 1080** | **99.26** | **99.23** | **99.98** | * | **99.75** | **99.98** | **100** |
|  | *-91.09* | *-89.18* | *-99.7* |  | *-95.9* | *-99.56* | *-99.88* |
| **Klebsiella pneumoniae KP ST11 OXA48** | **99.27** | **99.22** | **99.74** | **99.75** | * | **99.74** | **99.75** |
|  | *-91.73* | *-88.89* | *-94.46* | *-93.02* |  | *-94.3* | *-93.23* |
| **Klebsiella pneumoniae 440 1540** | **99.25** | **99.21** | **99.99** | **99.98** | **99.74** | * | **99.98** |
|  | *-90.4* | *-87.96* | *-99.8* | *-97.86* | *-95.59* |  | *-98.07* |
| **Klebsiella pneumoniae 646 1568** | **99.26** | **99.24** | **99.99** | **100** | **99.75** | **99.98** | * |
|  | *-90.94* | *-89.01* | *-99.77* | *-99.73* | *-95.95* | *-99.6* |  |
